# Biocompatible soft hydrogel lens as topical implants for diabetic retinopathy

**DOI:** 10.1101/2021.07.23.453466

**Authors:** Rajkumar Sadasivam, Gopinath Packirisamy, Mayank Goswami

## Abstract

Fashioned contact lenses can be converted into novel drug delivery vehicles. Ocular disease like diabetic retinopathy is a major microvascular complication where its early diagnosis and treatment is still a mystery to clinicians. Delivery of pharmaceuticals to the posterior part of eye is quite difficult, with exemption from the injectable formulations. Drug loaded hydrogel like contact lens implants can be utilized in place of commercial contact lens. Such hydrogel lens implants are developed with precise manner in which they are to be inserted as implants in the mice models. The present work is the preliminary outcome of our ongoing research work where the polymerized lens implant can be used as payload carrier for retinopathy. The aim is to propose such appropriate implants suitable for in vivo mice models.

## Introduction

Soft materials development in the form of contact lens implant for drug delivery of retinopathy is always challenging. Diabetic retinopathy (DR) is one of the most severe ocular complication where drug delivery to interior part is difficult. This is due to the presence of static barriers in cornea, blood and dynamic barriers like blood-retinal junction which inhibit the drug uptake into the ocular region. This inhibition causes lower bioavailability of drugs to the target1,2. Different types of therapeutic materials are developed for carrying out this task. Most of the ocular diseases of anterior region is treated with eye drops. However, the posterior ocular complications are treated with variety of materials such as hydrogels, implants, microneedles, solid lipid nanoparticles, dendrimers etc3,4.

These ocular carriers are administered through different routes such as topical, intravitreal, sub-conjunctival and subretinal. The former topical route is non-invasive and self-administered but has several disadvantages. The latter methods are injection based in which frequent procedure may damage the tissues. The drugs are washed off from anterior region due to corneal barrier. The other routes are also facing similar difficulties. Ocular implants which are topically administered are better option, but the choice of material is the crucial factor for design. Therefore, hydrogels in the form of therapeutic contact lenses are developed as topical implants. Polymeric hydrogels of natural or synthetic origin are widely used in drug delivery due to its intrinsic properties such as biocompatibility, biodegradability, increased drug resident time, sustained release etc. Despite the availability of effective injectable drug deliverables and aqueous pharmaceutical formulations for ocular delivery, stable hydrogels as contact lens implant is rarely developed. In recent times, Hydrogels have been emerged as promising drug delivery carriers due to their ability to meet the payload delivery challenges5,6. Hydrogels loaded with different combination of drugs are extensively used in the biomedical applications for drug delivery and tissue engineering. Few important necessary criteria for a better payload vehicle consists of a) Enhanced biocompatibility with appropriate selection of materials, b) protection of payload over prolonged time period of delivery, c) tunable release of payload from carrier and d) act as payload depot during therapy.

Transparent hydrogel is the foremost requirement for lens implant in mouse models followed by other characteristics such as hydrophilicity, high water content, durability, high drug loading capacity etc. These implants are prepared by various physical, chemical and physiochemical methods. Material properties of different soft implants suitable for ocular drug delivery along with its merits and demerits are studied7.

In the present work, Hydroxyethyl methacrylate and chitosan based hydrogels are developed with the aim of investigating the better choice of drug delivery carrier among the developed hydrogels for delivering the payloads. Therefore hydrogels of soft implants from above precursors were synthesized through different techniques. The suitability of ocular implant was confirmed via physical measurements. Preliminary studies such as biocompatibility, transmittance was performed. Based on the initial investigations, here we reported the appropriate or desired contact lens implant to be used for ocular drug delivery.

## Materials

Poly (2-hydroxyethyl methacrylate) (99.99%, Sigma), Triethylene glycol dimethacrylate (TEGDMA) (99.99%, Sigma), Chitosan (99.99%, Sigma), Ammonium persulfate (99%, SRL laboratories), Sodium metabisulphite (RANKEM laboratories), Gluteraldehyde, GA (99.99%, Sigma), 2-Hydroxyethyl methacrylate (99.99%, TCI), Ethylene glycol (SRL), tween-80 (SRL), acetic acid (97%, SRL Laboratories).

## Methods

### Preparation of PHEMA contact lens

Flexible soft hydrogel lens was prepared through free radical polymerization method. In brief, 5 mL of 2-Hydroxyethyl methacrylate (2-HEMA) monomer solution was added into a 15 mL glass vial and it has been mixed with 0.23 mL of TEGDMA which act as the cross-linking agent. This was added to the solvent mixture of DI water/ethylene glycol (1 mL/1.5 mL). After few minutes, 0.5 mL of 15 % sodium meta-bisulfite and 0.5 mL of 40 % of Ammonium persulfate was added and stirred for some stipulated time period. This act as the initiating agents of polymerization. The above polymeric mixture was poured into a polystyrene petri dish and kept overnight at room temperature for the polymerization reaction to occur. The obtained polymerized disc was removed and sliced into the required size using surgical knife and soaked in DI water for 4 h. The solvent has to be frequently changed in order to remove the unreacted monomer and the initiators. Finally the hydrogel discs are dried at room temperature for 48 h and stored for further use.

### Preparation of PHEMA hydrogel

Poly (2-hydroxyethyl methacrylate) soft hydrogel was prepared by simple homogenization method. 2% PHEMA (w/v) was added into 5 mL of DI water and continuously stirred for 48 h at 400 rpm. The hydrophobic PHEMA flakes absorbs water and swells to form stretchable uniform soft hydrogel and stored.

### Preparation of Chitosan hydrogel

Transparent aqueous stable chitosan hydrogels are prepared by chemical crosslinking method. In brief, 2% of chitosan is dissolved in 5 mL of DI water with 2% aqueous acetic acid. The above solution was stirred at room temperature for 24 h to get pale yellow colour. Then few drops of 0.5% tween-80 was added and filtered. Later 0.1% gluteraldehyde (1 mL) was added as crosslinking agent and again stirred for 30 min at room temperature. Further it was poured into a petri dish and dried at RT overnight to obtain cross-linked chitosan hydrogel. The hydrogel was dried at 45 °C for 12 h to remove the unreacted solvent and GA content.

### Transmissivity

The amount of light transmitted into the hydrogel lens was studied using UV-Visible spectrophotometer.

### Biocompatibility

HEK 293 (Human embryonic kidney) Cell line was procured from the cell repository of National Centre for Cell Science, India. Cells were maintained in Dulbecco’s modified Eagle’s medium (high-glucose) medium supplemented with 10% fetal bovine serum, 50 U/mL penicillin, 50 mg/mL streptomycin in a humidified atmosphere in 5% CO2 at 37 °C. 3-(4, 5-dimethylthiazol-2-yl)-2, 5-diphenyl tetrazolium bromide (MTT) assay was performed on animal cell lines using the procedures mentioned elsewhere to substantiate the biocompatibility of the different types of polymeric soft hydrogels such as PHEMA, HEMA and Ch.

### Microscopy analysis

Combined fluorescent staining dyes such as Rhodamine B and Hoechst 33342 was used to monitor the morphological deformation of cytoplasm and nucleus of mammalian kidney cells post treatment. Rhodamine B stains mitochondria and cytoplasm and it was visualized under GFP filter. Similarly, Hoechst 33342 dye stains the DNA that binds to the Adenosine-Thiamine (AT) pairs inside the nucleus through DAPI. Further, the mammalian cells treated with different soft materials are washed with PBS and stained with Rhodamine B of 1 μL (stock concentration of 1 mg/mL) for 10 min. The additional content of Rhodamine B has been removed post incubation period and added with PBS containing Hoechst 33342 dye (2 μL) (concentration of 10 mg/mL). PBS fixation was done on the above stained mammalian cells and the morphology of the cells were analyzed using fluorescence cell imaging microscopy (EVOS) through GFP and DAPI filter. The merged microscopic images represented the intact nucleus and cytoplasm of HEK 293 cells obtained from the combined filters.

## Results and Discussion

### Fabrication of hydrogel implants

Three different soft implant materials namely 2-HEMA, PHEMA and chitosan hydrogels are developed. 2-HEMA based hydrogel was found to have similar morphology to commercial contact lens (Fig 1A), whereas the PHEMA hydrogel appears as swollen gel with high water absorption (Fig1B). Visual observation of the gel found to have low transmittance and bulky nature which makes it undesirable as implant in mice model. Similarly, the chitosan based hydrogel was found to be very fragile and seems brittle in nature shown in Fig 1C. It was identified to be lesser desirable solid implant, however it is being used as effective injectable gels for ocular drug delivery applications8–10. The above observations are based on heuristic approach through physical examination of the hydrogels. In the end, the 2-HEMA based hydrogel was found to be appropriate implant as contact lens in mice models and thus selected for further studies. The selection is purely based on its applicability to mice eye and considered as suitable topical carrier.

**Fig 1.**
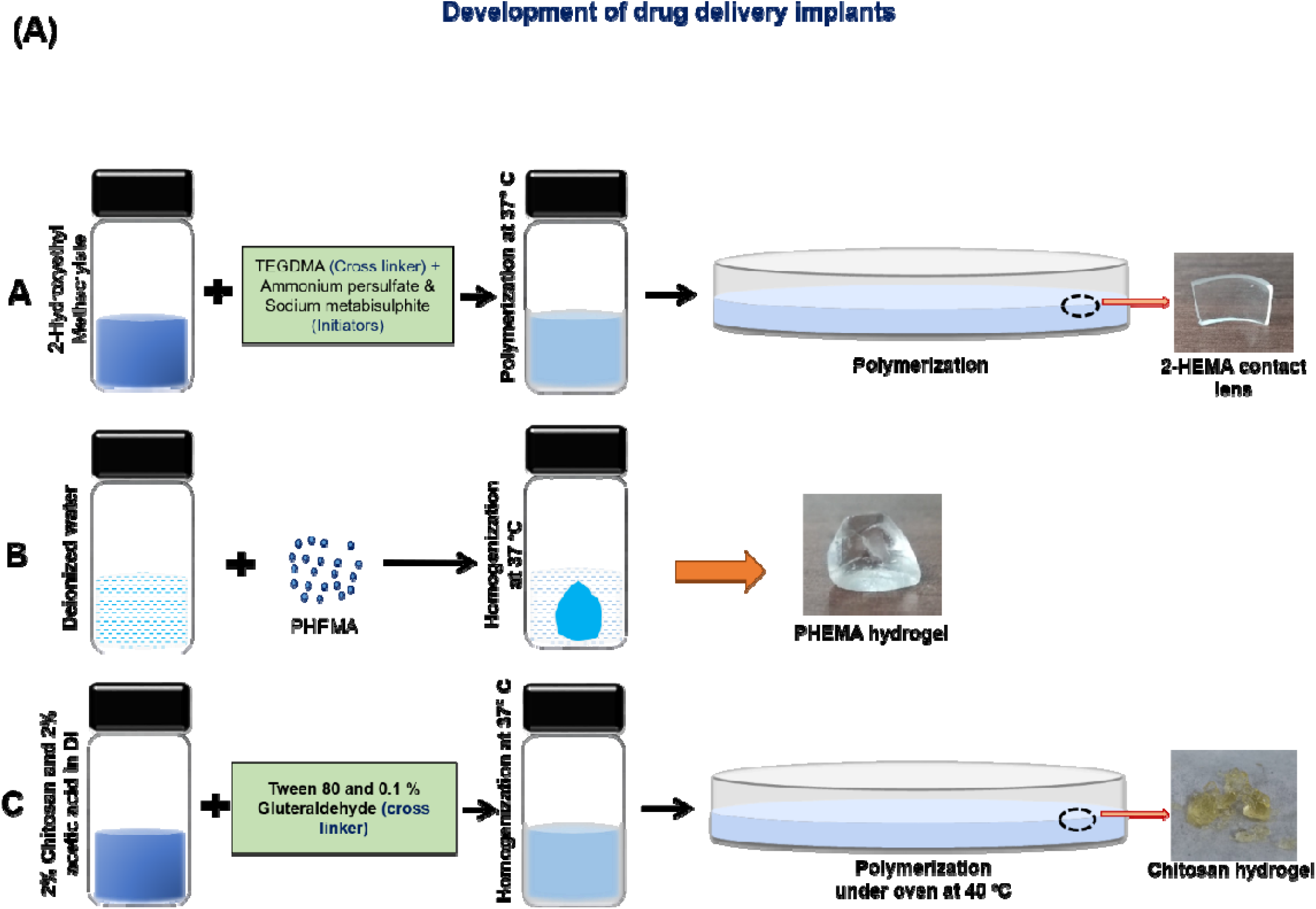
Development of polymeric soft material implants for drug delivery

### Biocompatibility

Soft materials developed are tested to evaluate its biocompatibility and also to select appropriate implant among different hydrogels for use. This study acted as prior checkpoints to avoid any toxic/immune reaction in mice eyes due to the use of hydrogels as ocular implants. The implants were sliced in circular discs according to the dimension of plates (30 mm cell culture plate) and cells were seeded over the material before incubation. Morphology of mammalian cells were examined through fluorescence microscopy. For this, Rhodamine B/Hoechst 33342 are used as staining dyes. Fig 2a-c represented the qualitative analysis of HEK 293 cells treated with three different hydrogels namely pHEMA, pHEMA lens and chitosan hydrogels. Hoechst 33342 stains the nucleus of the cell (visualized in DAPI filter) whereas the Rhodamine B stains the cytoplasm indicated over the periphery of the cells (shown in green filter). The morphology was observed to be intact even after 48 h of treatment in all three cases. Fig 2d represented the percentage viability of HEK 293 cell lines treated using hydrogels for 72 h and untreated control experiment was done for comparison. It was observed that all three soft hydrogels exhibited better biocompatibility (>99 % for 2-HEMA and PHEMA based lens and >85% for chitosan hydrogel). The percentage viability in treated sample is greater than 100% due to the proliferation of the cells during incubation. This implicated that the developed hydrogels act as suitable scaffold or substrate for the growth of mammalian cells. Furthermore, time dependent morphological structure of cells were evaluated. Fig 3a-c represented real time microscopic images of the HEK 293 cells treated with pHEMA hydrogel lens in the time intervals starting at 12 h, 24 h and 48 h, respectively. Henceforth, we confirmed that the developed hydrogels have excellent biocompatible properties and thus can be used as carriers for drug delivery. On the contrary, the physical property of chitosan and pHEMA hydrogels are undesirable for topical contact lens implant. Since the hydrogels are less translucent in nature which are verified through naked eye.

**Fig 2.**
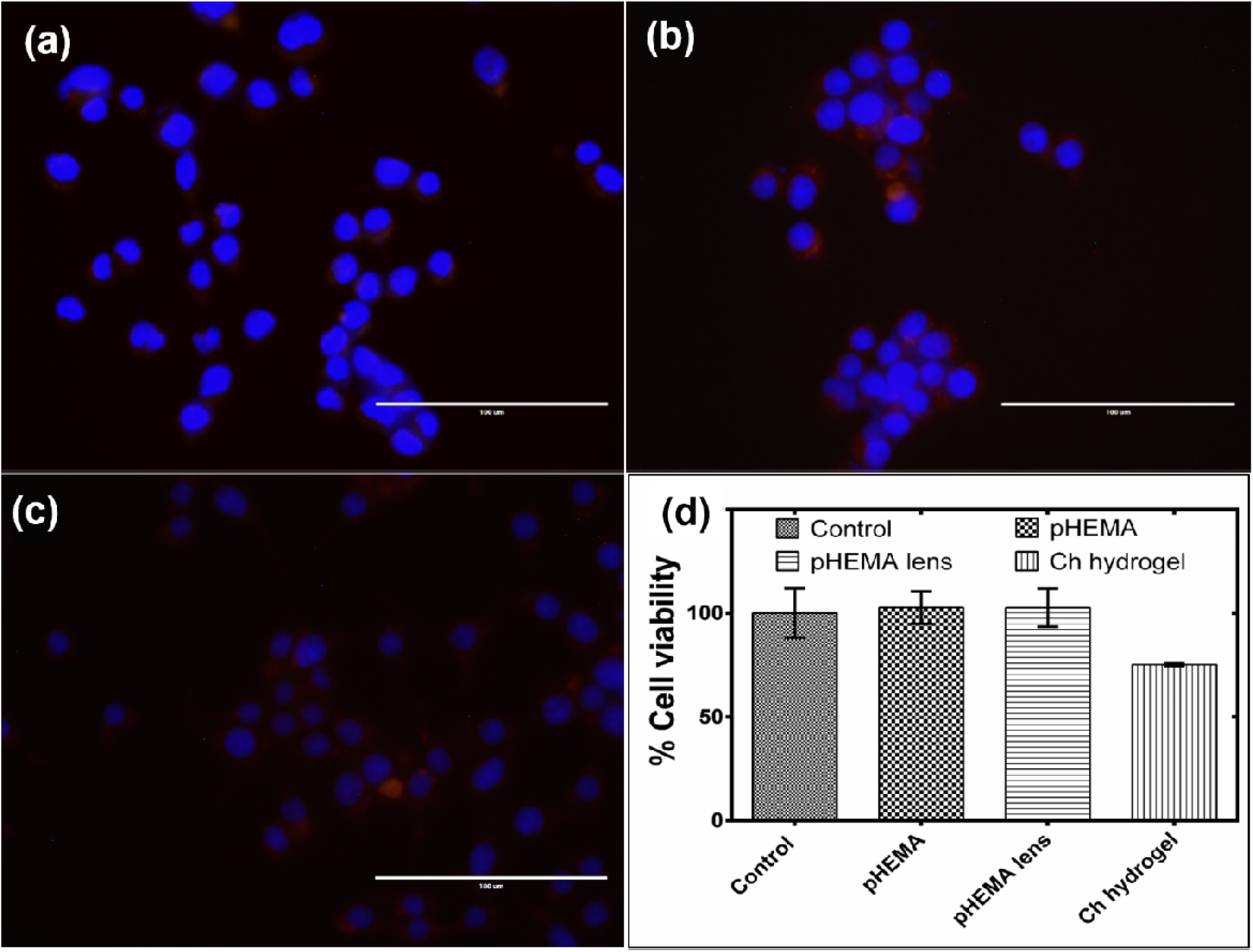
Biocompatibility studies of different soft hydrogel lenses after 48 h of treatment. (a-c) Qualitative microscopic analysis (*scale: 100 μm*) and (d) MTT assay (HEK 293 Human embryonic kidney cell lines was used for MTT assay)

**Fig 3.**
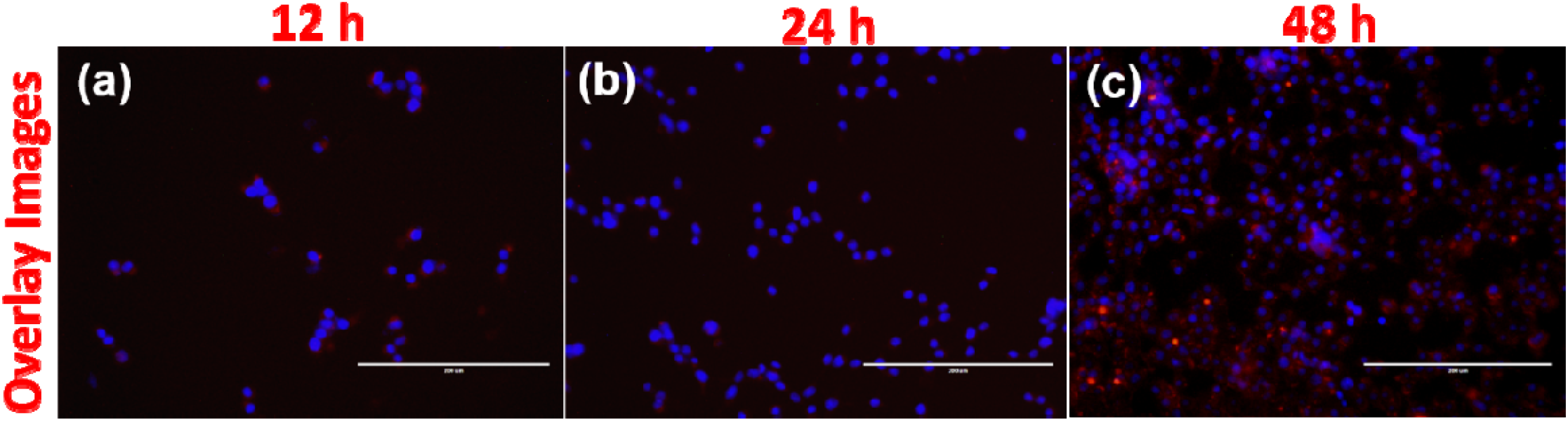
Time dependent Fluorescent microscopic images of HEK-293 cell lines treated with pHEMA hydrogel lens and stained with Hoechst 33342 (blue) and co-stained with Rhodamine B (red). a) 12 h, b) 24 h and c) 48 h. *Scale bar: 200 μm*.

### Transmissivity

The pHEMA hydrogel lens was visibly found to be highly transparent through naked eyes. Therefore, the transmissivity of pHEMA lens was confirmed through UV-Visible spectrophotometer. Fig 4 represented the percentage transmissivity of the lens with % transmittance on y-axis and wavelength (200-1000 nm) on x-axis. It was observed that the hydrogel lens exhibited continuous transmissivity in the visible region (400 – 700 nm), having maximum transparency ranging from 83.5 – 89.9%. This confirmed the applicability of hydrogel lens as implant for mice normal vision. The transparency and softness of implant lens can be further improved by reducing the thickness up to 1mm during its preparation.

**Fig 4.**
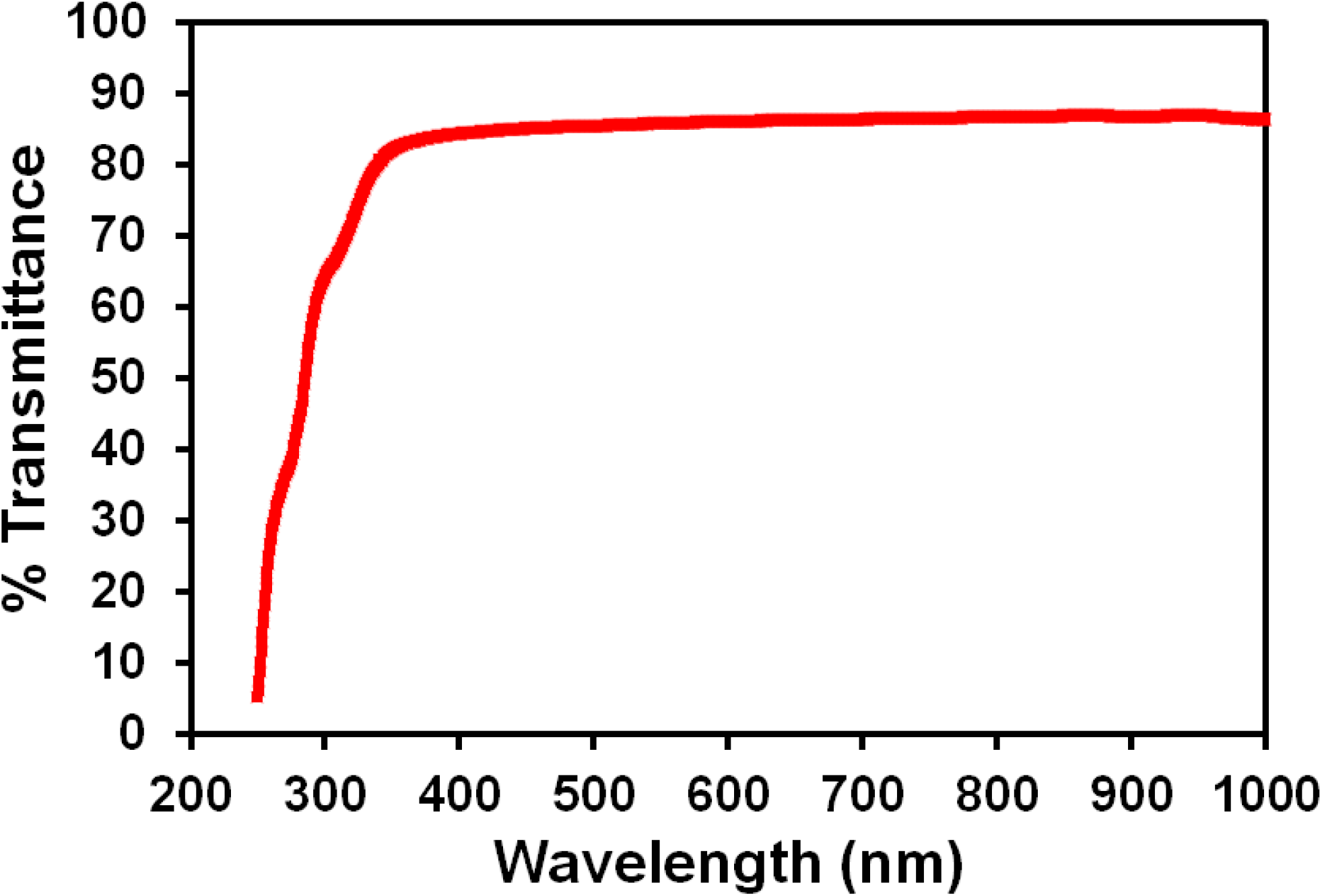
UV-Vis analysis showing % Transmissivity of pHEMA contact lens

## Summary

We have successfully developed soft hydrogel materials for ocular implants. Biocompatible properties are investigated quantitatively and qualitatively. Based on its applicability, the appropriate hydrogel lens was selected and further transmissivity was evaluated. In this work, the preliminary investigations confirmed the suitable carrier for topical ocular implant. Further supplementary studies need to be done before performing drug delivery of bevacizumab for retinopathy. Fig.5 represented the proposed utility of our developed hydrogel lens as implant. The following scanning scheme demonstrated the 3D image reconstruction of retina for our current research. Our future work involves real time imaging assisted diagnosis and therapy/treatment through ocular implants and pharmaceutical formulations.

**Fig 5.**
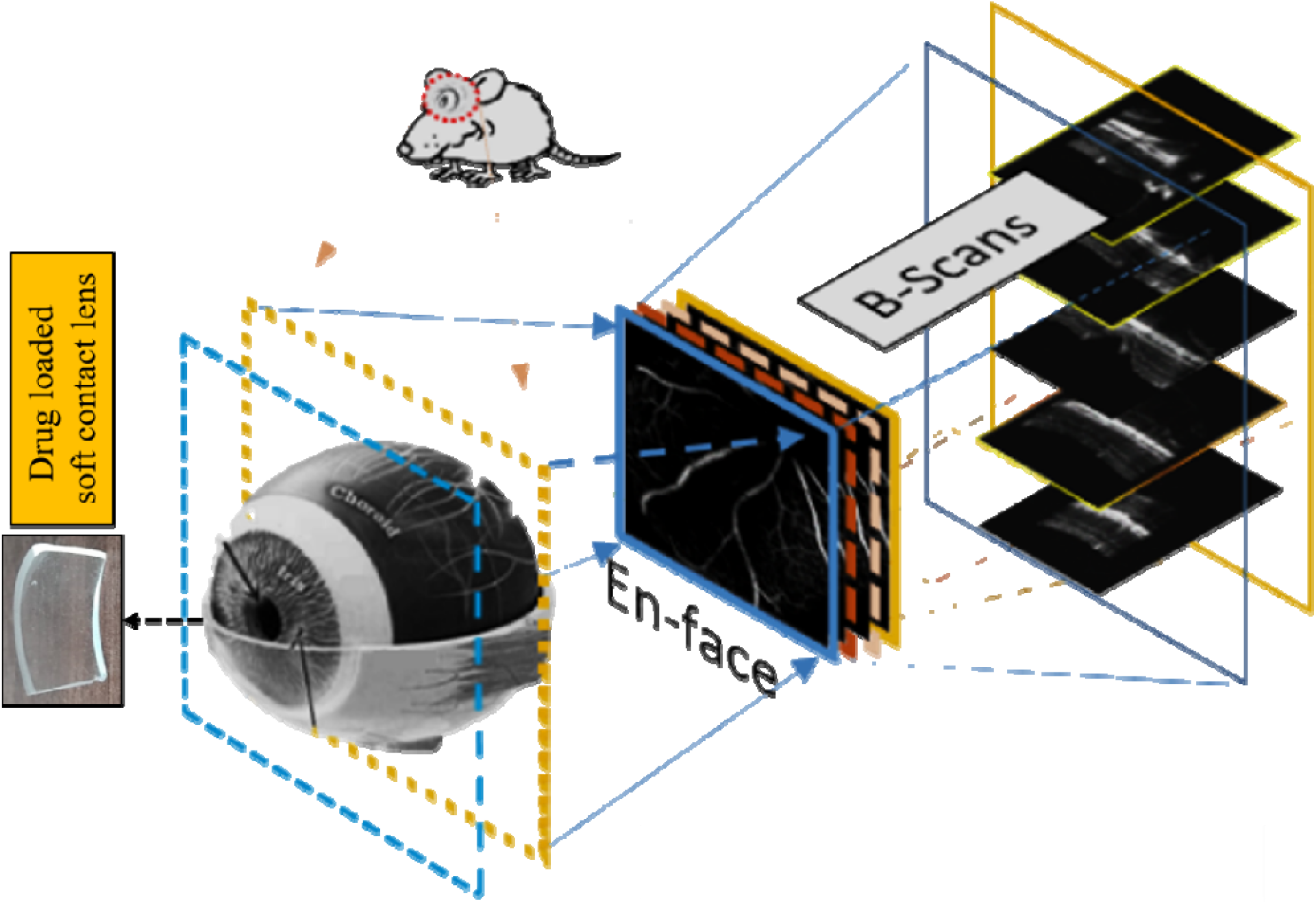
Proposed utility of the developed PHEMA hydrogel lens in mouse eye and its scanning scheme. (Dr. Mayank’s lab^©^)

## Acknowledgements

Dr. Rajkumar Sadasivam is thankful to DST-SERB for the research fellowship and sincere thanks to the Department of Physics and Department of Bioscience and Bioengineering of the Indian Institute of Technology Roorkee. Prof. P Gopinath (Co-Principle Investigator) is highly acknowledged for bringing the implant idea in ocular drug delivery. This work is financially supported by the Department of Science and Technology-Science and Engineering Research Board (DST-SERB) under IMPRINT-2 scheme, Project grant code: IMP/2018/001045, Government of India.

## Note

Authors declare no competing interest

